# Systemic TLR7 agonism moderates disease severity in acute DSS-induced colitis in mice

**DOI:** 10.64898/2026.11.07.765085

**Authors:** Daniel Kazimierczyk, Brian Seed, Nitya Jain

## Abstract

Toll-like receptor 7 (TLR7) is an endosomal sensor of single-stranded RNA, studied for its antiviral and pro-inflammatory functions. Its role in intestinal inflammation is undefined, and no approved or clinical-stage therapy for inflammatory bowel disease (IBD) engages this pathway. Here we report that systemic administration of the reference TLR7 agonist GS-9620 (vesatolimod) moderates disease severity in acute dextran sodium sulfate (DSS)-induced colitis in mice. GS-9620-treated mice lost less weight, had lower Disease Activity Index scores, and were protected from DSS-induced colon shortening. Immunophenotyping of the spleen showed reduced Th17 and Tr17 populations and an increased proportion of regulatory T cells. These findings indicate that pharmacological TLR7 agonism is not merely tolerated in the inflamed intestine but is associated with a shift of the systemic immune compartment toward a regulatory phenotype. The results identify TLR7 as a candidate disease-moderating target in IBD and motivate direct tests of causality, route and chronic-disease relevance.

## Introduction

Toll-like receptor 7 (TLR7) is an endosomal sentinel primarily expressed by plasmacytoid dendritic cells (pDCs), macrophages, and B cells, where it detects single-stranded RNA(1). Upon ligand binding, TLR7 triggers Type I Interferon (IFN) signaling essential for orchestrating innate and adaptive immune responses(2). TLR7 signaling downstream of endosomal activation branches into two distinct arms(2). The MyD88-dependent arm activates NF-κB and drives production of pro-inflammatory cytokines. A second arm, IRF7 activation, drives Type I IFN production(3). While IRF7-dependent Type I IFN production is classically described in plasmacytoid dendritic cells, TLR7-driven Type I IFN induction has also been demonstrated in macrophages(4). Type I IFNs induced downstream of TLR7 have been linked to intestinal epithelial barrier integrity and homeostatic balance(5, 6). In the gastrointestinal tract, these signaling cascades maintain a critical balance between luminal surveillance and mucosal homeostasis.

A small but consistent literature indicates that engaging TLR7 in the intestine is protective rather than injurious. Depleting the enteric virome with antiviral treatment worsens dextran sodium sulfate (DSS) colitis and reconstitution with TLR3 or TLR7 agonists or with inactivated rotavirus ameliorates disease(7). Mice lacking both TLR3 and TLR7 are more susceptible to experimental colitis, and combined TLR3 and TLR7 variants influence the severity of ulcerative colitis in patients(7). Consistent with this, the TLR7 agonist imiquimod given systemically reduced disease activity, histological damage, colonic myeloperoxidase activity and circulating TNF-α and IL-6 in DSS colitis(8), and attenuated TNBS colitis while inducing colonic IFN-β and TGF-β, raising IL-10, and driving CCR9-dependent accumulation of Foxp3^+^ Treg cells in the inflamed colon(9). In addition to reprogramming immunity, TLR7 engagement also acts on the epithelial barrier. Intracolonic imiquimod enhanced barrier function in rats, lowering epithelial conductance and macromolecular passage(10). An orally administered PEGylated TLR7 ligand prevented intestinal barrier disruption and bacterial translocation while inducing Reg3β, Reg3γ and IL-22, with *Tlr7*-deficient animals showing the opposite phenotype(11). In contrast, topical imiquimod applied to skin induces psoriasiform dermatitis and secondarily exacerbates DSS colitis through dysbiosis and altered colonic macrophage and B cell compartments, an effect transferable with feces(12, 13).

Two mechanisms plausibly account for a therapeutic effect. The first is the output of the IRF7 arm, which can be engaged without dominant MyD88/NF-κB signaling. Type I IFN acts protectively in the gastrointestinal tract, imposing an antiproliferative state that supports epithelial differentiation, maturation and barrier function(5). The locus containing *IFNAR* has been identified as a susceptibility region for human IBD(14). Interestingly, this appears to be a property of tonic rather than acute interferon: mice lacking STAT1 spontaneously develop intestinal inflammation accompanied by dysbiosis and expansion of Th17 cells, and blocking IL-17 signaling prevents the downstream myeloid pathology(15). Type I IFN also constrains Th17 development directly, acting through innate-cell IL-27, such that TRIF- or IFNAR-deficient animals develop more severe autoimmune inflammation with elevated IL-17(16). The second mechanism is innate tolerance. Repeated low-dose stimulation with the TLR7 agonist 1V136 rendered myeloid cells progressively hyporesponsive and induced cross-tolerance to TLR2, TLR7 and TLR9 ligands. This limited disease in experimental autoimmune encephalomyelitis and antibody-mediated arthritis, acting through upregulation of IRAK-M and SHIP-1(17). Repeat imiquimod exposure similarly induces the negative regulators A20/TNFAIP3 and IRAK3 alongside IL-10 and TGF-β1, in a macrophage MMP10-dependent manner(18), and persistent dosing with R848 produces tachyphylaxis of both behavioral and transcriptional inflammatory responses, with cross-refractoriness to LPS(19). Under an intermittent dosing schedule, a TLR7 agonist is hypothesized establish a regulated state marked by negative regulators of TLR signaling, IL-10 and with Type I IFN output preserved. Whether that state is reachable in the inflamed intestine, and whether it translates into reduced disease, is the question this study addresses.

GS-9620 (vesatolimod) is a selective, orally bioavailable TLR7 agonist that has been characterized pharmacologically and advanced into clinical trials for chronic hepatitis B and HIV, with a well-defined dosing and safety profile in animals and humans(20-23). It therefore provides a well-controlled reference agent to test whether pharmacological TLR7 agonism aggravates or moderates disease in an acute model of intestinal injury and inflammation. Here we report that systemic GS-9620 reduces clinical and gross-anatomical measures of disease severity in a DSS-induced model of acute colitis that is associated with a shift in the colonic T cell compartment away from Th17 and toward regulatory phenotypes. These data identify TLR7 as a candidate disease-moderating target in intestinal inflammation.

## Results

### Systemic GS-9620 reduces clinical disease severity in acute DSS colitis

To test whether pharmacological TLR7 agonism modifies the course of acute intestinal inflammation, adult male C57BL/6 mice received 3% (w/v) DSS in drinking water for 7 days and were then returned to normal drinking water until day 10. GS-9620 (3 mg/kg) or solvent was administered by intraperitoneal injection on days 2, 4 and 6, during the induction phase of disease (**Figure 1A**). Animals were assessed daily and tissues were collected at the peak of disease (day 7) and after three days of recovery (day 10).

**Figure 1:**
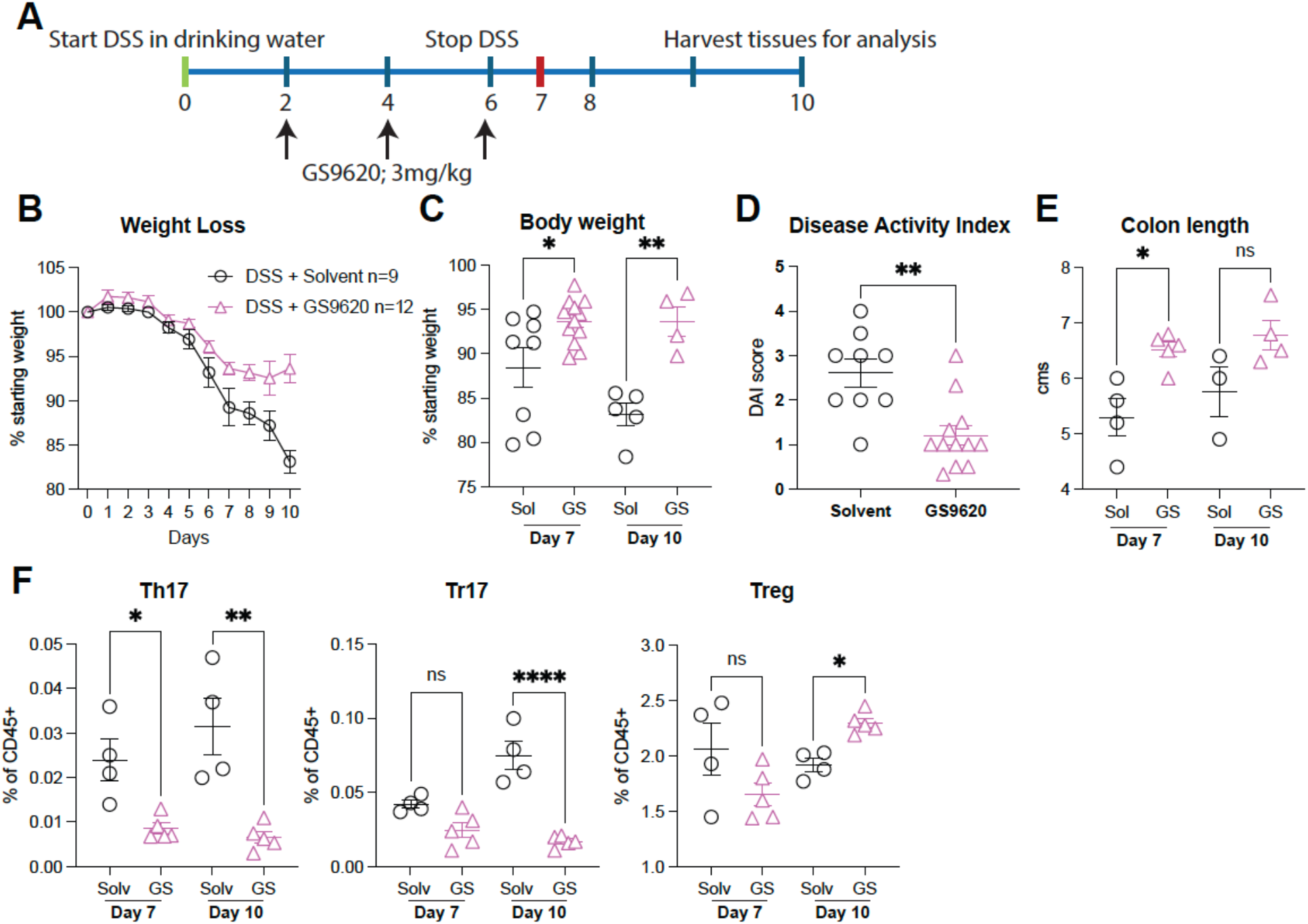
TLR7 agonism moderates acute DSS induced colitis in mice. A. Adult male C57BL/6 mice were given 3% (w/v) DSS in drinking water for 7 days, after which they were placed on normal water till Day 10. GS-9620 (3mg/kg) was administered by intra-peritoneal injection on days 2, 4, and 6. **B, C**. Body weights were measured daily. **D**. DAI score was calculated as described here(24). **E**. At necropsy on Days 7 and 10, the colon from the cecal junction to the rectum was removed and its length recorded. **F**. Single cell suspensions prepared by enzymatic digestion of the colonic tissue were stained with antibodies to identify FOXP3^+^ Tregs, RORgt^+^ Th17 and FOXP3^+^RORgt^+^ Tr17 cells. Data are representative of 2 independent experiments. Error bars are standard error mean. Two-tailed, unpaired t-test was used to calculate significance. * *p* <0.05, ** < 0.01, *** < 0.001, **** <0.0001.

Mice receiving GS-9620 were protected from DSS-induced weight loss. While both groups lost weight from approximately day 5 onward, the GS-9620-treated group diverged from solvent-treated controls during the second half of DSS exposure and remained above controls through the recovery period (**Figure 1B**). The difference reached significance at both harvest points, with GS-9620-treated animals retaining 93.6% versus 88.4% of starting body weight at day 7 (*p* < 0.05) and 93.6% versus 83.1% at day 10 (*p* < 0.01) (**Figure 1C**). Notably, solvent-treated mice continued to lose weight after DSS withdrawal, whereas GS-9620-treated mice recovered, suggesting an effect on resolution as well as on peak injury.

Disease Activity Index (DAI), a composite of weight loss, stool consistency and rectal bleeding(24), was substantially lower in GS-9620-treated mice than in controls (**Figure 1D**; *p* < 0.01), with most treated animals scoring at or below 1.5 while control scores clustered between 2 and 4.

Colon shortening, a gross-anatomical correlate of colitis severity, was likewise attenuated. At day 7, colons from GS-9620-treated mice were significantly longer than those from solvent-treated controls (**Figure 1E**; *p* < 0.05). At day 10 the same direction of effect was present but did not reach significance, reflecting partial recovery of colon length in the control group during the washout period.

### TLR7 agonism shifts splenic immune cell compartments toward a regulatory phenotype

To determine whether the clinical benefit was accompanied by changes in the immune compartment, single-cell suspensions of spleens were analyzed by flow cytometry (**Figure 1F**). The spleen was chosen for three reasons. First, DSS colitis is a systemic illness rather than a purely local one, and splenic cellularity and immune composition change measurably over the disease course(25). Second, since GS-9620 was administered intraperitoneally, splenocytes were used to represent the systemic readout of this route of exposure. Third, lamina propria leukocytes at peak disease consist largely of infiltrating cells, so any treatment that reduces mucosal damage will mechanically reduce the infiltrate, and a lower Th17 frequency in colonic tissue cannot readily be separated from simply having less colitis. A shift measured in the spleen is not subject to that confound.

GS-9620 treatment reduced the proportion of RORγt^+^Th17 cells among CD45^+^ cells at both day 7 (*p* < 0.05) and day 10 (*p* < 0.01). FOXP3^+^RORγt^+^ Tr17 cells were unchanged at day 7 but markedly reduced at day 10 (*p* < 0.0001). Conversely, FOXP3^+^ regulatory T cells were comparable between groups at day 7 and significantly increased in GS-9620-treated animals at day 10 (*p* < 0.05).

Taken together, these data show that systemic TLR7 agonism during the induction phase of DSS colitis reduces clinical and gross-anatomical disease severity and is associated with a shift of the systemic T cell compartment away from Th17 and toward regulatory phenotypes.

## Discussion

TLR7 is a pro-inflammatory innate immune sensor, and agonizing it systemically during active intestinal injury would be predicted to worsen colitis. However, across clinical, gross-anatomical and immunological measures, mice treated with GS-9620 had less severe disease than solvent-treated controls, and the effect persisted through the recovery phase.

TLR7 signaling bifurcates between a MyD88/NF-κB arm that drives inflammatory cytokine production and an IRF7 arm that drives Type I IFN. Type I IFN signaling has been linked to intestinal epithelial barrier integrity and to restraint of Th17 responses, and both the reduction in RORγt^+^ cells at peak disease and the later expansion of FOXP3^+^ regulatory T cells are consistent with that axis.

Our study has several limitations. First, DSS colitis is a chemically induced epithelial injury model. It is reproducible and well validated for assessing mucosal damage and repair, but it does not reproduce the chronic, relapsing, immune-driven pathology of human ulcerative colitis. Protection in this setting establishes that TLR7 agonism can dampen acute mucosal injury; it does not establish efficacy in chronic disease, which requires relapsing models. Second, GS-9620 was given intraperitoneally, producing systemic exposure. Sustained systemic Type I IFN induction carries a recognized liability profile, including influenza-like symptoms and, with chronic exposure, autoimmune phenomena. Any therapeutic application of this biology would need to establish whether a local interferon signature can be achieved without systemic interferon exposure. Finally, *Tlr7* is X-linked and escapes X-inactivation in a fraction of female immune cells, and sex differences in TLR7 responsiveness are documented(26, 27). The exclusive use of male animals here is a meaningful limitation, and sex should be treated as a biological variable in follow-up work.

## Methods

### Animals

Animal studies were conducted under protocols approved by the Institutional Animal Care and Use Committee of Massachusetts General Hospital. Male C57BL/6 (Stock No. 00664), were purchased from the Jackson Laboratory and housed in Specific Pathogen Free (SPF) facilities at MGH for the duration of the experiments.

### Colitis induction and treatment

Colitis was induced with 3% (w/v) dextran sodium sulfate (DSS; MP Biomedicals, colitis grade, mol. wt. 36,000-50,000) in drinking water for 7 days, after which animals were returned to normal drinking water until day 10. GS-9620 (Vesatolimod; Selleckchem Cat.No.S7221) was formulated in 5% DMSO, 10% Solutol HS15 and 85% saline (0.9% NaCl), and administered by intraperitoneal injection at 3 mg/kg on days 2, 4 and 6. Control animals received an equal volume of solvent on the same schedule.

### Clinical assessment

Body weight was recorded daily and is expressed as a percentage of starting weight. Disease Activity Index was scored as previously described(24) combining weight loss, stool consistency and the presence of occult or gross rectal bleeding.

### Tissue collection

At necropsy on day 7 or day 10, the colon was removed from the cecal junction to the rectum and its length measured.

### Flow cytometry

Single cell suspensions were prepared from the spleens of adult animals following mechanical homogenization. RBCs were lysed using ACK lysis buffer. A maximum of 3 x 10^6^ cells were stained with fluorophore-conjugated antibodies. After surface staining, cells were fixed and permeabilized using the Foxp3/Transcription Factor Staining Buffer Set (Thermofisher Scientific) and stained with intracellular transcription factor antibodies. Dead cells were excluded using LIVE/DEAD Fixable Blue Dead Cell Stain Kit (Invitrogen Life Technologies). Samples were acquired on a 5 laser Cytek Aurora spectral analyzer. Single color controls were made using UltraComp eBeads for spectral unmixing on the cytometer.

### Statistics

Data are presented as individual animals with mean ± SEM. Comparisons between solvent and GS-9620 groups at each timepoint were made using a two-tailed, unpaired *t*-test. Significance thresholds: \**p* < 0.05, \*\**p* < 0.01, \*\*\**p* < 0.001, \*\*\*\**p* < 0.0001. Analyses were performed in GraphPad Prism. Data shown are representative of two independent experiments.

## Acknowledgments

We are thankful for the resources of the Center for Computational and Integrative Biology at MGH that supported this study.

## Author contributions

DK, NJ performed experiments and analyzed data. NJ and BS designed experiments and NJ wrote the paper with input from all authors.

